# Quantitative accuracy and precision in multiplexed single-cell proteomics

**DOI:** 10.1101/2021.09.03.458853

**Authors:** Claudia Ctortecka, Karel Stejskal, Gabriela Krššáková, Sasha Mendjan, Karl Mechtler

**Affiliations:** Research Institute of Molecular Pathology (IMP), Vienna BioCenter (VBC), Campus-Vienna-Biocenter 1, 1030 Vienna, Austria; Institute of Molecular Biotechnology of the Austrian Academy of Sciences (IMBA), Vienna BioCenter (VBC), Dr. Bohr-Gasse 3, 1030 Vienna, Austria; The Gregor Mendel Institute of Molecular Plant Biology of the Austrian Academy of Sciences (GMI), Vienna BioCenter (VBC), Dr. Bohr-Gasse 3, 1030 Vienna, Austria

**Author notes:** Correspondence: Claudia Ctortecka, Karl Mechtler, Research Institute of Molecular Pathology (IMP), Campus Vienna Biocenter 1, 1030 Vienna, Austria.

## Abstract

Single-cell proteomics workflows have considerably improved in sensitivity and reproducibility to characterize yet unknown biological phenomena. With the emergence of multiplexed single-cell proteomics, studies increasingly present single-cell measurements in conjunction with an abundant congruent carrier to improve precursor selection and enhance identifications. While these extreme carrier spikes are often >100-times more abundant than the investigated samples, undoubtedly the total ion current increases, but quantitative accuracy possibly is affected. We here focus on narrowly titrated carrier spikes (i.e., <20x) and assess their elimination for comparable sensitivity at superior accuracy. We find that subtle changes in the carrier ratio can severely impact measurement variability and describe alternative multiplexing strategies to evaluate data quality. Lastly, we demonstrate elevated replicate overlap while preserving acquisition throughput at improved quantitative accuracy with DIA-TMT and discuss optimized experimental designs for multiplexed proteomics of trace samples. This comprehensive benchmarking gives an overview of currently available techniques and guides conceptualizing the optimal single-cell proteomics experiment.

## Introduction

Single-cell proteomics has been demonstrated as a viable complement to single-cell transcriptomics studies with striking sensitivity. Those single-cell analyses of presumed homogenous cell populations have attributed biological variability at both the transcriptome and the proteome levels.^1,2^ Previously, most protein analyses with single-cell resolution have been antibody-based or were limited to large cells such as oocytes.^3,4^ More recent technological innovations now allow for the hypothesis-free proteome analysis of single mammalian cells.^5–7^ First of such aimed at overcoming the limited sensitivity of available mass spectrometers through isobaric labeling.^3,8^ Isobaric labels use their identical mass with different isobaric distribution, allowing the simultaneous analysis of multiple samples and their quantification upon fragmentation within one MS^n^ scan. SCoPE-MS (Single Cell ProtEomics by Mass Spectrometry) combines TMT-multiplexed single cells with a 200-cell congruent carrier sample.^8^ The highly abundant carrier overcomes adsorptive losses before MS analysis, boosts the peptide signals during MS1 scans, therefore, increasing the signal to noise ratio (S/N) of the peptide precursor and serves fragment ions for identification. Following the initial publication, such congruent carrier spikes were employed to improve the triggering of peptides of interest at varying ratios from 25x up to 500x.^7,9,10^

While an abundant carrier improves peptide identifications by increasing ion counts, selecting appropriate acquisition parameters and carrier compositions are crucial to preserving quantitative accuracy.^11–13^ Such imbalanced levels of multiplexed carrier samples with ratios of 200 or higher were demonstrated to possibly impact biological conclusions.^9,11,13,14^ The effects of extreme ratios on ion suppression^15^, ratio compression^16,17^ and quantification accuracy^9^ were previously described for standard and trace samples. The latter was addressed by increasing the number of ions sampled from each precursor^12,18,19^, constrained by the injection time (IT) and the automatic gain control (AGC) target. However, increasing the total number of ions sampled per precursor capitalizes the ions originating from the carrier within imbalanced samples.^13,20^ Additionally, the thereof lengthened cycle times reduce the number of MS/MS scans within one analytical run, inflating missing data between replicates due to precursor stochasticity.^21^

Recently, Cheung and colleagues evaluated the ‘carrier proteome’ effects for trace samples proposing a maximum level of a congruent carrier (~20x) for optimal ion statistics and quantification accuracy.^13^ They and others thoroughly discuss the need for appropriate MS acquisition parameters and S/N filtering when performing MS-based single-cell proteomics experiments.^12,13^ However, it remains unclear which levels of excess carrier provide the optimal balance between sensitivity and accuracy or whether even drastically reduced ratios impair quantitative precision. Therefore, we demonstrate the applicability and confirm the need for S/N filtering to improve data quality in various multiplexed experimental designs. Moreover, with alternative multiplexing or acquisition strategies, we discuss the impact of precursor co-isolation and precursor stochasticity in the analysis of trace samples. This study aims to outline the advantages of available experimental setups and compile critical aspects, including identification rates, measurement accuracy, acquisition variability, and missing quantitative data in single-cell proteomics experiments.

## Methods

### Sample preparation

Cells were pelleted, washed with phosphate-buffered saline (PBS) and stored at −80°C until they were lysed using a methanol:chloroform:water solution (4:1:3), sonicated and dried to completeness in a speed-vac concentrator. The dry protein pellets were resuspended in 8M urea in 10mM HCl. Prior to alkylation with iodoacetamide (40mM, 30min at room temperature (RT)), the samples were adjusted to 200 mM Tris/HCl pH 8.0 and reduced using dithiothreitol (50mM, 37°C, 30min). The reduced and alkylated samples were diluted to a final concentration of 4M urea in 100mM Tris/HCl pH 8 and digested with LysC (Wako, enzyme:protein 1:100) for 3 hours at 37°C, if indicated. Tryptic samples were subsequently diluted to 2M urea in 100mM Tris/HCl pH 8 and digested over night at 37°C (Promega, enzyme:protein 1:100). Samples were adjusted to pH 2 using 10% trifluoroacetic acid (TFA) and desalted using C18 solid-phase extraction cartridges (SPE, C18 Sep-pak, 200 mg Waters) eluted with 40% acetonitrile (ACN) in 0.1% TFA. SPE eluate volume was reduced using a vacuum centrifuge and labeled according to manufacturer’s instructions. Briefly, samples were labeled in 100mM TEAB and 10% ACN for 1 hour at RT. Unreacted TMT reagent was quenched with 5% hydroxylamine/HCl for 20 minutes at RT and subsequently mixed corresponding to each sample pool.

### LC MS/MS analysis

Samples were measured on a Orbitrap Exploris 480 Mass Spectrometer (Thermo Fisher Scientific) with a reversed phase Dionex UltiMate 3000 high-performance liquid chromatography (HPLC) RSLCnano system coupled via a Nanospray Flex ion-source equipped with FAIMS Pro (Thermo Fisher Scientific), which was operated at a constant compensation voltage of −50 V. Chromatographic separation was performed on nanoEase M/Z Peptide BEH C18 column (130Å, 1.7μm, 75μm × 150mm, Waters, Germany) developing a two-step solvent gradient ranging from 1.2 to 30% over 90 min and 30 to 48% ACN in 0.08% formic acid within 20min, at a flow rate of 250nL/min.

SCoPE-MS and SCoPE2 acquisition strategies were performed as published, with small adaptations. For SCoPE-MS and SCoPE2 samples briefly, full MS data were acquired in the range of 395–1,800 or 450-1,600m/z at a resolution of 60,000, respectively. The maximum AGC was set to 3e6 and automatic IT. Top 20 or 7 multiply charged precursor ions (2-3 or 2-4) with a minimum intensity of 2e4 were isolated for higher-energy collisional dissociation (HCD) MS/MS using a 1 or 0.7 with 0.3 Th offset isolation window, respectively. Precursors were accumulated until they either reached an AGC target of 1e5, or a maximum IT of 250 or 300ms, respectively. MS/MS data were generated with a normalized collision energy (%NCE) of 34, at a resolution of 60,000, with the first mass fixed to 110m/z. Upon first fragmentation precursor ions were dynamically excluded (dynEx) for 20 or 30 seconds, respectively.

Full MS data of multiplexed carrier experiments were acquired in a range of 375-1,200m/z with a maximum AGC target of 3e6 and automatic IT at 120,000 resolution. Top 10 multiply charged precursors (2-5) over a minimum intensity of 5e3 were isolated using a 0.7 Th isolation window and acquired at a resolution of 60,000 at a fixed first mass of 110m/z with a maximum AGC target of 1e5 or IT of 118ms and dynamically excluded for 120 seconds. TMT10-plex (TMT10) precursors were fragmented at an NCE of 34 and TMTpro at an NCE of 32.

TMT_zero_ experiments acquired similarly but precursors were selected using a ‘targeted mass-difference method’ for 3 seconds cycle time. For this, the delta masses of 5.0105 Da or 10.0209 Da were used to select only precursors with a matching partner intensity to the most intense one, with a mass tolerance of 10ppm. Targeted precursors were isolated with a 0.7 Th isolation window at an AGC target of 1e5, for maximum 250ms IT. Selected precursors were acquired at a resolution of 45,000 and subsequently excluded with for 100seconds.

DIA-TMT experiments were acquired in the range of 400–800m/z at a resolution of 45,000. The AGC was set to 2e5 with automatic maximum IT and a 5 Th isolation windows including 1Th overlap with the first mass fixed to 120m/z. This corresponds to 80 DIA windows with a cycle time of 6 seconds. TMT10 samples were fragmented with a stepped NCE of 30, 37.5, 45 and TMTpro with 25, 30, 40.

### Data analysis

RI quantification was performed within the Proteome Discoverer environment (version 2.3.0.484) using the in-house developed, freely available PD node “IMP-Hyperplex” (https://pd-nodes.org) to extract the intensity and S/N values for all a RIs at a mass tolerance of 10ppm. Quality control of raw data was performed using RawTools.^22^ Venn diagrams were generated using BioVenn.^23^

Peptide identification was performed using the standard parameters in SpectroMine™ 2.0 against the human reference proteome sequence database (UniProt; version: 2018-11-26 accessed April 2019). Briefly, we performed a specific tryptic search with maximum two missed cleavages limiting peptides to 7-52 amino acids. We included carbamidomethylation on cysteins, TMT or TMTpro on lysine and all N-terms as fixed modifications and acetylation on protein N-terms and methionine oxidation variable. By default, SpectroMine™ performs ideal mass tolerance calculations at MS and MS/MS levels and mass calibration for each feature. Subsequently, identifications were filtered for 1% FDR on PSM, peptide and protein-group level for further processing.

For SCoPE-MS and SCoPE2 re-analysis, raw files were obtained from the following repositories and processed as indicated above (MSV000084660, MSV000083945 MSV000082077).^7,8^

TMT spectral libraries were generated from the DDA files with above indicated parameters using a customized script provided by Oliver Bernhard from Biognosys (available on GitHub: ctorteckac/DIA-TMT).^24^ Libraries were searched with Spectronaut™ performing mass tolerance calculations and spectra matching based on extensive mass calibration. The most intense peak within the previously defined mass tolerance is then selected and matched with a minimum of three matching fragment ions per MS/MS scan. RT alignments are based on iRT Reference Strategy with minimum R^2^ 0.8. ‘Mutated’ decoys with scrambled sequences are filtered for 1% FDR on precursor and protein levels.

## Results and discussion

To directly compare diverse multiplexing strategies for ultra-low input samples, we performed labeling of HeLa digest in bulk. We combined 150pg peptide input per TMT-label, which we will from now on refer to as ‘single cell’, with carrier titrations. Based on previous findings concerning accurate ratio reporting, we performed TMT10 experiments with a maximum carrier of 10x.^16^ As similar studies for the sixteen channel TMTpro are lacking, but several studies demonstrated quantitative implications of a >20x carrier, we extended this titration to 20x.^9,13^ Additionally, we evaluated a ‘dual carrier’ with the carrier distributed equally across two TMT channels to reduce extreme ratios but still boost ions. To overcome isobaric interference of the carrier sample, we did not include adjacent channels in the quantitative evaluations. For SCoPE-MS and SCoPE2 (SCoPE) experiments, we adapted the respective acquisition parameters and experimental setup detailed in the methods section (Fig. 1a).^7,8^ SCoPE2 is described to yield about 1,000 proteins from real single-cell measurements.^7^ Our reanalysis of their raw data identified up to 700 protein groups, while adopting their experimental setup, we reproducibly identify ~1,300 proteins from diluted bulk samples and are therefore confident to reflect previously published protocols accurately (Fig. 1b; Supplemental Fig. 1a).

**Figure 1:**
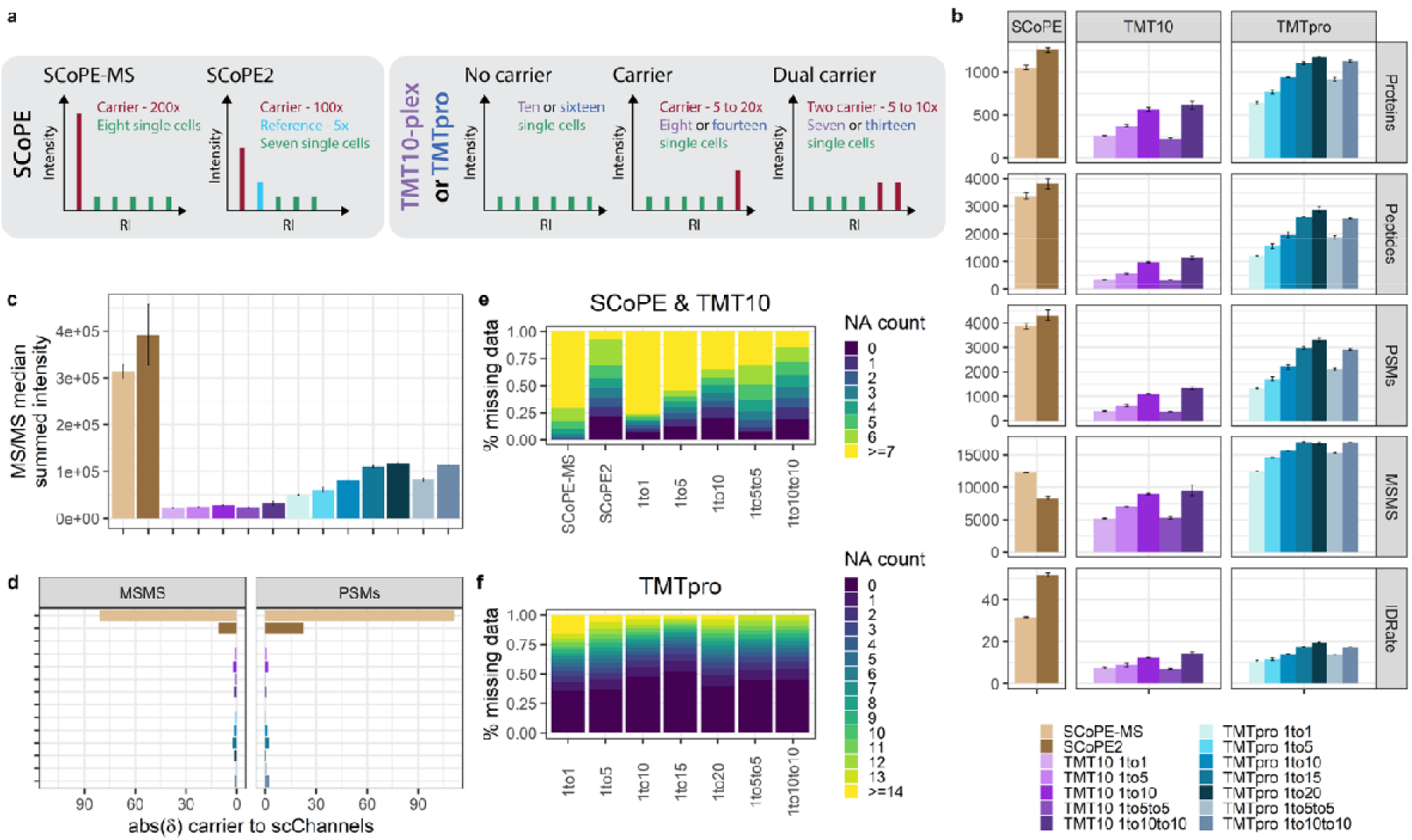
Characterization of TMT multiplexed carrier titrations. **(a)** Graphical illustration of experimental setups and carrier compositions. **(b)** Identified proteins, peptide groups, PSMs, number of MS/MS scans, ID-rates, **(c)** median summed MS/MS intensity and **(d)** delta between expected and acquired carrier to ‘single cell’ ratio across all MS/MS scans or PSMs for SCoPE (brown), TMT10 (purple) and TMTpro (blue) samples at indicated carrier spikes. Median and median absolute deviation (mad) is shown. Percent missing quantitative data in **(e)** SCoPE, TMT10 and **(f)** TMTpro carrier titrations per PSM.

### Abundant carrier spikes enhance protein identifications but suffer from ratio compression

We identified around 1,000 proteins for SCoPE and TMTpro 15-20x samples, in detail SCoPE-MS experiments with a 200x carrier and 250ms max IT yielded 30% more MS/MS scans than SCoPE2 with 100x carrier and 300ms max. IT (Fig. 1b). Nevertheless, the 50% identification rate (ID-rate) of SCoPE2 outperformed all other experimental setups presented in this study, similarly to their published raw files (i.e. 30% ID-rate) from real single-cell measurements (Supplemental Fig. 1a).^7^ TMTpro 10-20x samples triggered over 16,000 low intensity MS/MS scans with only 20% ID-rate and finally 15% less protein identifications compared to SCoPE2. Likewise, the 60% increased peptide amount of respective TMTpro and TMT10 samples yields more intense MS/MS scans and superior ID-rates (Fig. 1b). Additionally, the larger TMTpro reagents (419 Da) require only 30 %NCE compared to 34% for the TMT10 tag (344 Da). Surprisingly, although we injected similar peptide amounts, average MS/MS scans and ID-rates declined in split-carrier experiments for TMT10 and TMTpro. We speculate, that one abundant carrier relatively contributes more productive signal and less noise compared to two lower ones. Concluding, due to the increased peptide amount per injection, extreme carrier spikes result in high-intensity MS/MS spectra but only in conjunction with increased AGC targets and max. IT serves the necessary fragment ions for enhanced protein identifications (Fig. 1b-c).

To investigate the quantitative accuracy of multiplexed measurements, we first included all MS/MS scans with reporter ion (RI) signals and determined the delta of expected to acquired RI intensities. Strikingly, we observed that the ‘single cell’ signal in SCoPE-MS experiments was compressed by close to 50%, while SCoPE2 drastically improved ratio compression to only 10% (Fig. 1d). Similarly, in published SCoPE datasets we observe severe signal compression for SCoPE-MS and only to a lesser extent in SCoPE2 (Supplemental Fig. 1b). Next, we only considered identified scans and observed that the most abundant MS/MS scans provide sufficient fragment ions for identification but exhibit strong ratio compression. Further, only ~50% of all MS/MS were within the carrier range (+/− 50% of expected ratio) for SCoPE experiments (Fig. 1d; Supplemental Fig. 2). This is in stark contrast to most balanced TMT10 and TMTpro experiments with more than 80% of all MS/MS scans within the expected carrier ratio (Supplemental Fig. 2). We speculate that high-intensity MS/MS spectra exhibit elevated noise levels, which impacts measurement accuracy in the presence of an abundant carrier and is aggravated in real-single cell samples.

Considering the abundant carrier spike within the RI cluster, we dissected intensity profiles of individual channels and observed isobaric interference in SCoPE experiments, as expected (Supplemental Fig. 3a-b; Supplemental Fig. 4a-b). Therefore, they exclude the adjacent channel for single cells but establish an empty or reference channel for quality control and normalization.^7,8,14^ Lower carrier ratios did not exhibit isobaric interference, but adjacent channels were nevertheless excluded from subsequent analysis (Supplemental Fig. 3c-f). To investigate whether measurement variation and signal intensity parallels, we correlated the coefficient of variation (CV) between ‘single cell’ RIs within one MS/MS scan to the average RI S/N. For this, we combined three technical replicates, removed all MS/MS scans with only carrier or missing RIs and determined the %CV. In our experimental setup, all ‘single cell’ channels are distributed equimolar, theoretically resulting in 0% CV. As expected, most MS/MS scans with low S/N and multiple missing quantitative values show high variance. Despite enhanced average MS/MS intensity in SCoPE experiments, the mean ‘single cell’ S/N is lower than balanced TMT10 and TMTpro experiments (Fig. 2a-f). While SCoPE-MS experiments present a decreased median of 25% CV compared to 30% in SCoPE2, the latter indicates a trend towards small %CV in high S/N MS/MS scans (Fig. 2a-b). Further, the 200x carrier in SCoPE-MS leads to a detrimental suppression of the ‘single cell’ RIs giving rise to almost 75% missing data. In contrast, the reduced 100x carrier yielded higher ‘single cell’ S/N, resulting in over 90 or 75% of all MS/MS scans with at least one or two RIs, respectively (Fig. 1e). This, however, disagrees with the re-analysis of published single-cell data, where SCoPE-MS presents 4-times elevated single cell RI S/N compared to SCoPE2 with almost no missing quantitative data across all PSMs (Supplemental Fig. 4c-e).

**Figure 2:**
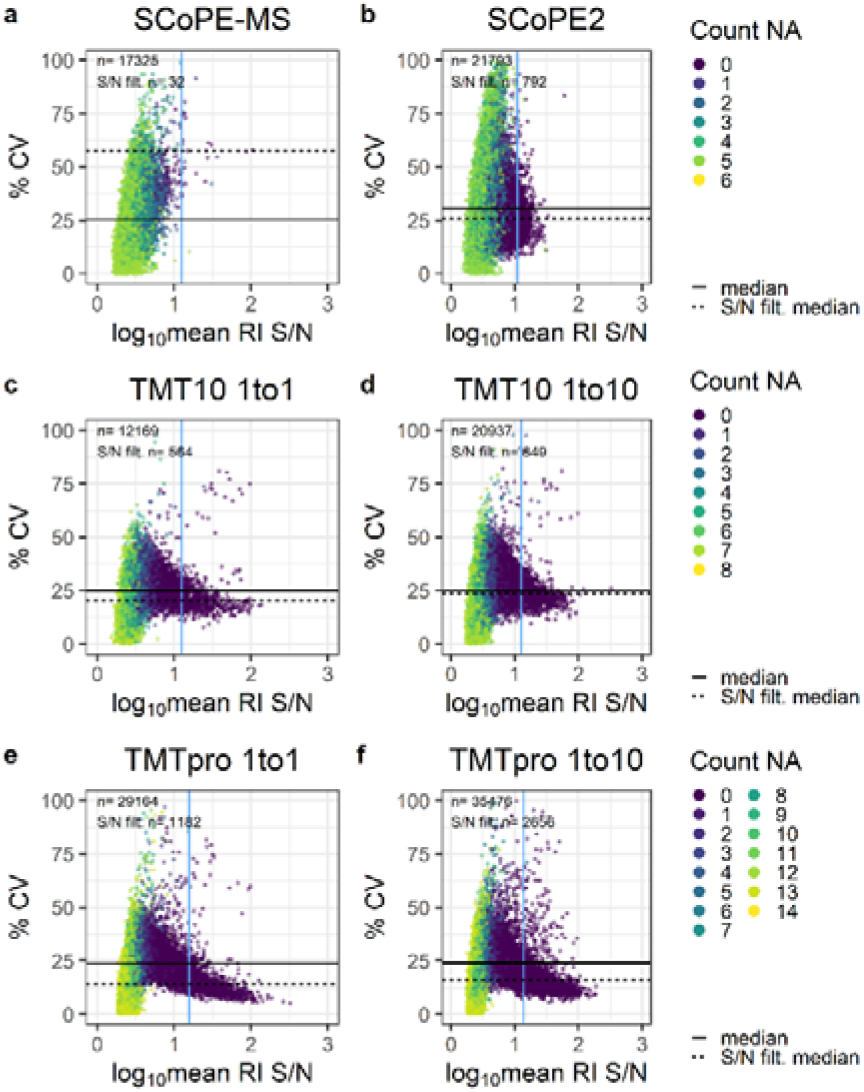
Quantification accuracy of various carrier titrations. Percent CV across ‘single cell’ channels and log_10_ mean RI S/N for **(a-b)** SCoPE, **(c-d)** TMT10 and **(e-f)** TMTpro samples at given carrier ratios. The horizontal solid and dashed lines indicate median S/N across all MS/MS scans or post-S/N filtering, respectively. The vertical blue line specifies the S/N filter cut-off. Colors reflect the number of missing ‘single cell’ RIs per MS/MS scan.

Based on analogous observations, quality controlling via RI S/N filtering was introduced with *SCPCompanion*,^13^ which we applied to our datasets, reducing the number of MS/MS scans and removing almost all scans with missing RIs. In detail, a minimum RI S/N of 12.6 for SCoPE2 eliminated over 96% of all MS/MS scans but improved the median CV by 5%. Similarly, for TMTpro no-carrier and 10x samples over 90% of all MS/MS scans were removed, but median CV was enhanced by 10%. In most experimental setups but especially across the limited carrier TMTpro samples, high RI S/N MS/MS scans trend toward low %CV (Fig. 2a-f). This was not observed in the re-analysis of published SCoPE datasets, which we attribute mainly to biological and technical variance. Nevertheless, it is noteworthy that quantitative confidence of real single-cell proteomics samples benefits from RI S/N filtering (Supplemental Fig. 4c-d). While identifications and measurement stability of bulk diluted TMTpro >10x and SCoPE2 experiments are comparable, ratio compression and quantitative inaccuracy in the latter suggests limiting the carrier to a maximum of 20x in combination with appropriate S/N filters (Fig. 1b, d; Fig. 2b-f; Supplemental Fig. 2; Supplemental Fig. 3 b-f).

### An alternative labeling strategy reveals frequent precursor co-isolation

Based on these findings, we aimed to preserve the advantages of an abundant carrier but remove the extreme ratio from the RI cluster for improved quantification accuracy. For this, in anticipation that the targeted quantitation of only ‘single cell’ derived peptides would greatly reduce the impact of inter-channel ratio compression, we made use of the defined mass difference provided by differential labeling of carrier and sample peptides with TMT_zero_ (224.152 amu) and TMT10 (229.162 amu) reagent, respectively. We digested the samples with Lys-C to label peptides on the C- and the N-termini, increasing the mass difference between the carrier and the ‘single cell’ channels.^25^ We combine TMT10 labeled ‘single cell’ peptide input with an abundant TMT_zero_ carrier at varying ratios starting with an equivalent of ten TMT10 labeled cells (i.e., 1to1) up to 200 times the combined ‘single cell’ peptide input (i.e., 1to20; Fig. 3a). Emanating from the mass separation, TMT_zero_ labeled carrier precursors highlight TMT10 labeled ions with identical characteristics. Therefore, ‘single cell’ precursors are selected for fragmentation despite being close to or below detection limit, theoretically without impairing ‘single cell’ quantification.

**Figure 3:**
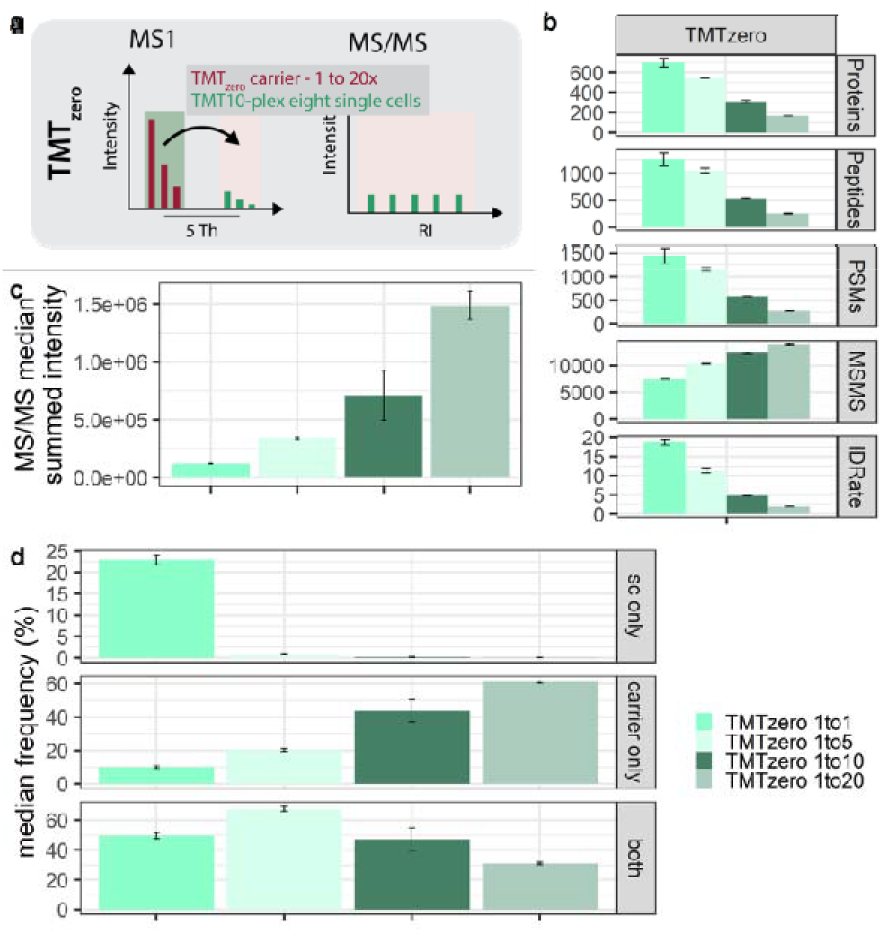
Measurement variability with inter-carrier spikes using TMT_zero_. **(a)** Graphical illustration of the TMT_zero_ triggering strategy. **(b)** Protein groups, peptide groups, PSMs, MS/MS scans, ID rates, **(c)** median summed MS/MS intensity and **(d)** percent median frequency of MS/MS scans across ‘single cell’ RIs (sc only), only carrier RI (carrier only) and co-isolation of carrier and ‘single cell’ precursors within one MS/MS scan (both) at indicated carrier spikes. Median and mad are shown.

Like inter-channel experiments, an abundant TMT_zero_ carrier repeatedly increased high-intensity MS/MS scans, but protein identifications declined with elevated carrier ratios (Fig. 3b-c). Within the TMT_zero_ approach, stemming from the different mass of the TMT10 and the TMT_zero_ tag, discrimination of ‘single cell’ versus carrier identifications is feasible. The 126-channel (i.e., fragment mass of TMT_zero_) was therefore excluded to overcome isobaric interference of mixed spectra and allowing to estimate co-isolation of a carrier precursor by the presence of a RI signal with 126.128 Da. This enables to estimate the frequency of only ‘single cell’, carrier or mixed MS/MS scans across the carrier titration. Interestingly, the relative frequency of co-isolating ‘single cell’ and carrier precursor increases with a 5× but decreases at ≥10x carrier ratio (Fig. 3d). Based on this, we speculate that the 20% reduced ID-rate across increasing TMT_zero_ carrier is partly due to the reduced number of TMT10 precursors, owing to the mass-based segregation of carrier and sample peptide ion species (Fig. 3b). Further, we observed that extreme congruent carrier spikes increase the chance of isolating only a carrier precursor 5-fold compared to balanced experiments (Fig. 3d). These findings indicate that in conjunction with an abundant carrier, it is likely that most PSMs correspond to a carrier-derived identification rather than the single cells. Consequently, the carrier must equally represent all single-cell precursors for accurate acquisition, which could be challenging for heterogeneous samples.

Despite low protein identifications, we evaluated measurement stability and quantification accuracy of the TMT_zero_ experimental setups. Corroborating earlier observations with similar inter-channel carrier ratios, we observed stable ‘single cell’ signal but elevated isobaric interference in the 126 and adjacent channels (Supplemental Fig. 5a-c). While the median CV below 25% across all MS/MS scans in TMT_zero_ 1× experiments decreased to only 13% after S/N filtering, a ≥10x carrier spike resulted in frequent missing values and up to 30% CV (Supplemental Fig. 5d-f). We conclude that removing the carrier from the multiplexed ‘single cells’ via TMT_zero_ elevates ‘single cell’ RI S/N but at extreme ratios impairs protein identifications and measurement accuracy (Fig. 3b, Supplemental Fig. 5d-f). Moreover, the mass difference between ‘single cells’ and carrier revealed close to 50% co-isolation and up to 60% carrier-only quantitative data in imbalanced ultra-low input samples (Fig. 3d). This suggests that a congruent carrier indeed improves MS/MS triggering and serves fragment ions; however, the identical features do not discern between solely a carrier or a ‘single cell’ PSM. Consequently, we speculate that all multiplexed ultra-low input experiments suffer a similar frequency of co-isolation and convoluted RI clusters.

### Intentional co-isolation reduces missing data in ultra-low input samples

TMT_zero_ experiments allowed to estimate unintentional co-isolation, RI convolution and its impact on MS/MS-based quantification accuracy (Fig. 3c-d; Supplemental Fig. 5a-f), as previously discussed by many.^13,17,25–31^ Additionally, we and others found detrimental amounts of missing data in multi-batch data-dependent proteomics experiments (Fig. 1e-f).^21,24,32–34^ This is most prominently addressed via data-independent acquisition (DIA), which our group recently extended to multiplexed samples.^24^ While co-isolation is non-negotiable with our 5 Th DIA-TMT method (i.e., in contrast to 0.7 Th in standard DDA), the prescheduled acquisition strategy theoretically generates no-missing data across multiple analytical runs (Fig. 4a). In detail, our small window DIA-TMT method allows to uniformly generate abstract 3D maps, comprised of RT, precursor m/z and RI intensity. These 3D maps or ‘proteome signatures’ entail convoluted RI quantification of a reproducible set of *in bona fide* precursors across all analytical runs. While we intentionally co-isolate multiple precursors to expedite sampling and provide consistent ‘proteome signatures’, convoluted RIs distinguish cell types down to single protein knock-outs.^24^ In contrast, the stochastic nature of data-dependent acquisition (DDA) methods, especially in analyzing ultra-low input samples, generates detrimental amounts of missing data. Consequently, this demands most quantitative data be computationally generated across large sample cohorts. However, the obvious application of any single-cell technology to characterize tissues or cellular subpopulations requires quantitative profiles of hundreds or thousands of samples, which is facilitated via our sensitive DIA-TMT strategy.^24^

**Figure 4:**
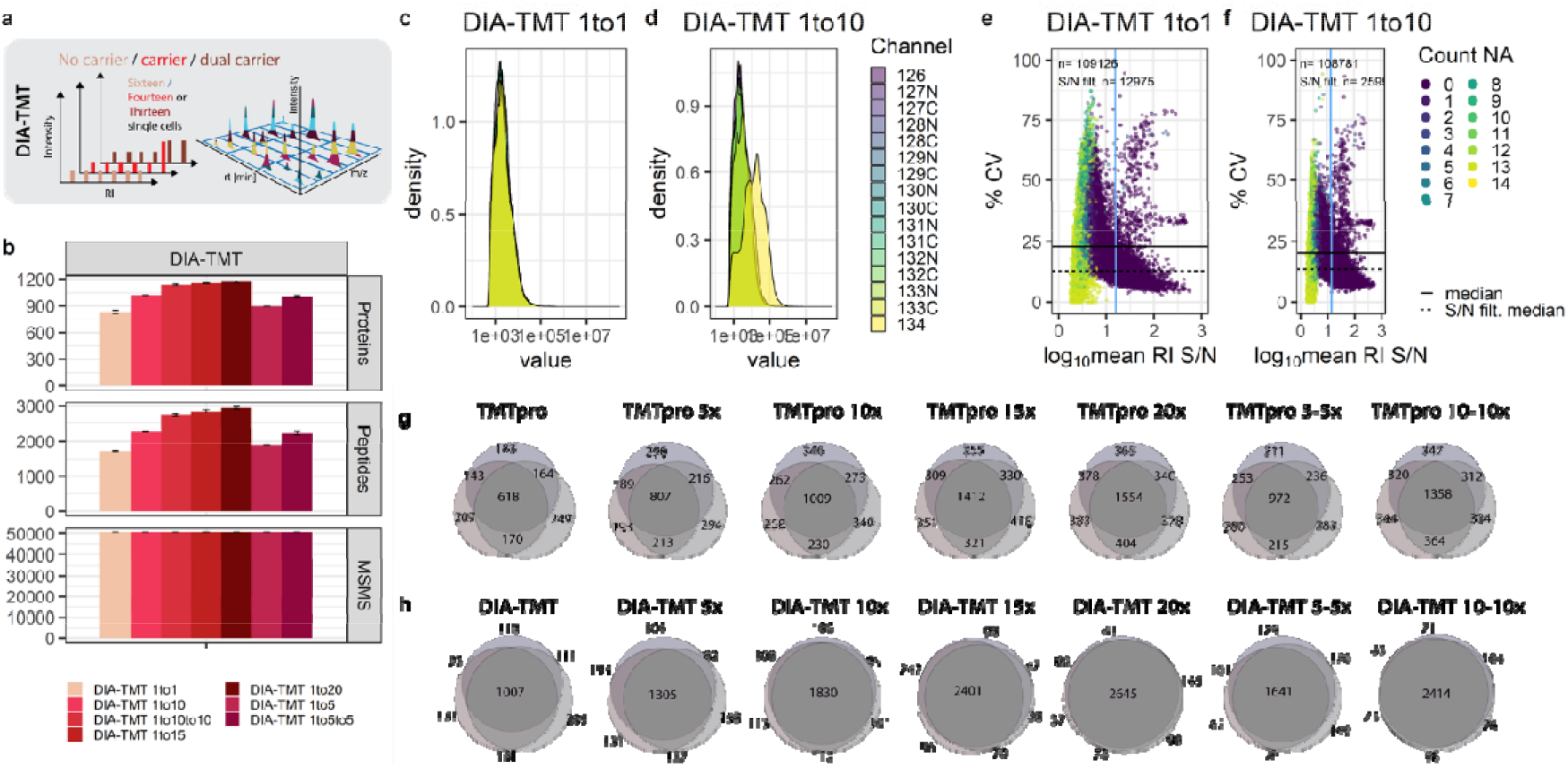
Impact of intentional co-isolation on ‘single cell’ variability and accuracy. **(a)** Graphical illustration of DIA-TMT acquisition strategies with carrier titrations. **(b)** Protein groups, peptide groups and MS/MS scans of DIA-TMT samples at indicated carrier spikes. Median and mad is shown. **(c-d)** RI intensity distributions across all MS/MS scans and **(e-f)** %CV against log_10_ mean RI S/N for DIA-TMT samples at indicated carrier spikes. The horizontal solid and dashed line indicate median S/N across all MS/MS scans or post-S/N filtering, respectively. The vertical blue line indicates the S/N filter cut-off. Colors indicate the number of missing single-cell RIs. Replicate overlap of **(g)** DDA and corresponding **(h)** DIA-TMT samples based on unique peptides.

Accordingly, we evaluated the quantification accuracy and reproducibility of DIA-TMT in conjunction with the TMTpro carrier titrations. Interestingly, the DIA-TMT samples yielded slightly increased protein identifications at similar PSMs to corresponding DDA measurements (Fig. 1b; Fig 4b). We speculate that this is because of the decreased cycle time and optimized fragmentation due to the stepped collision energy providing optimal fragmentation for co-isolated precursors with different charge states. Further, the measurement variance of no-carrier samples is comparable to DDA experiments but increases in combination with a congruent carrier spike (Fig. 4c-d; Supplemental Fig. 3e). While we observed similar overall RI intensities for DDA and DIA measurements, median RI S/N increased by 40% for the latter (Fig. 2 e-f; Fig. 4 e-f).

Interestingly, despite some distorted scans at high RI S/N, the DIA-TMT strategy presented close to 20% CV between ‘single cells’ across all carrier titrations. As previously discussed, S/N filtering further decreased the median CV to around 10%, corresponding to the lowest ‘single cell’ variation across all experiments (Fig. 4e-f). However, to constitute the complete ‘proteome signature’ the convoluted RI cluster attributes quantification to a set of co-isolated precursors rather than a single peptide species. Lastly, to directly compare replicate overlaps in DDA and DIA, we intersected unique peptide identifications. DIA-TMT increased replicate overlap by 25% in contrast to corresponding DDA samples across all carrier titrations (Fig. 4g-h). With low measurement variance, exceptional accuracy and close to 90% replicate overlap, DIA-TMT demonstrates its potential to overcome missing data at comparable quantitative accuracy in ultra-low input samples.

## Conclusion

We dissect different multiplexing strategies at extensive carrier titrations to investigate the impact on identification rates, reproducibility, quantification accuracy and measurement interference. Interestingly, we observe almost a linear increase in protein identifications across the low carrier titrations for both isobaric reagents. Moreover, we find that congruent carrier spikes, effectively contribute ions to the ‘single cells’ and consequently increase MS/MS intensities. Already a small carrier (<20x) improves ID-rates, which eventually plateaus at ≥100x ratio for 60-minute gradients. The SCoPE2 acquisition parameters and the 50% carrier decrease compared to SCoPE-MS, reduced ion suppression, increased ID rates, and improved measurement accuracy (Fig. 1b, d; Fig. 2a-b; Fig. 5a-d; Supplemental Fig. 2). While even lower carrier spikes on average resulted in less intense MS/MS scans and lower ID-rates, with less extreme RI ratios we observed no ratio compression, less measurement variability, and predominantly fewer missing values (Fig. 1b-f; Fig. 2c-f; Fig. 5a-d). However, even non-stringent S/N filtering often eliminated around 90% of MS/MS scans, suggesting that the RI S/N of such ultra-low input samples is suboptimal across experimental setups (Fig. 2c-f). This is especially concerning as diluted bulk digests utilized in this study contain less chemical background than real single-cell samples.^13,35^ Expectedly, *SCPCompanion* advised RI S/N filtering prior to database searching dramatically reduces protein identifications across all conditions and highlights the carrier limitation of <20x (Fig. 5b).^13^ Interestingly, the carrier abundance in TMT_zero_ samples parallels with the frequency of carrier co-isolation and therefore decreasing protein identifications. Despite that, for TMT_zero_ 50% more protein groups surpassed RI S/N filtering compared to all other experimental setups (Fig. 5a-b). We therefore speculate, that our current TMT_zero_ acquisition strategy despite being highly accurate and selective, suffers from inefficient triggering, which could be improved with more stringent precursor selection. Further, the quantitative accuracy of S/N filtered MS/MS scans indicates that a real-time search MS3-based approach to offset-trigger solely if the carrier precursor is identified would benefit the method (Supplemental Fig. 5d-f).^36,37^

**Figure 5:**
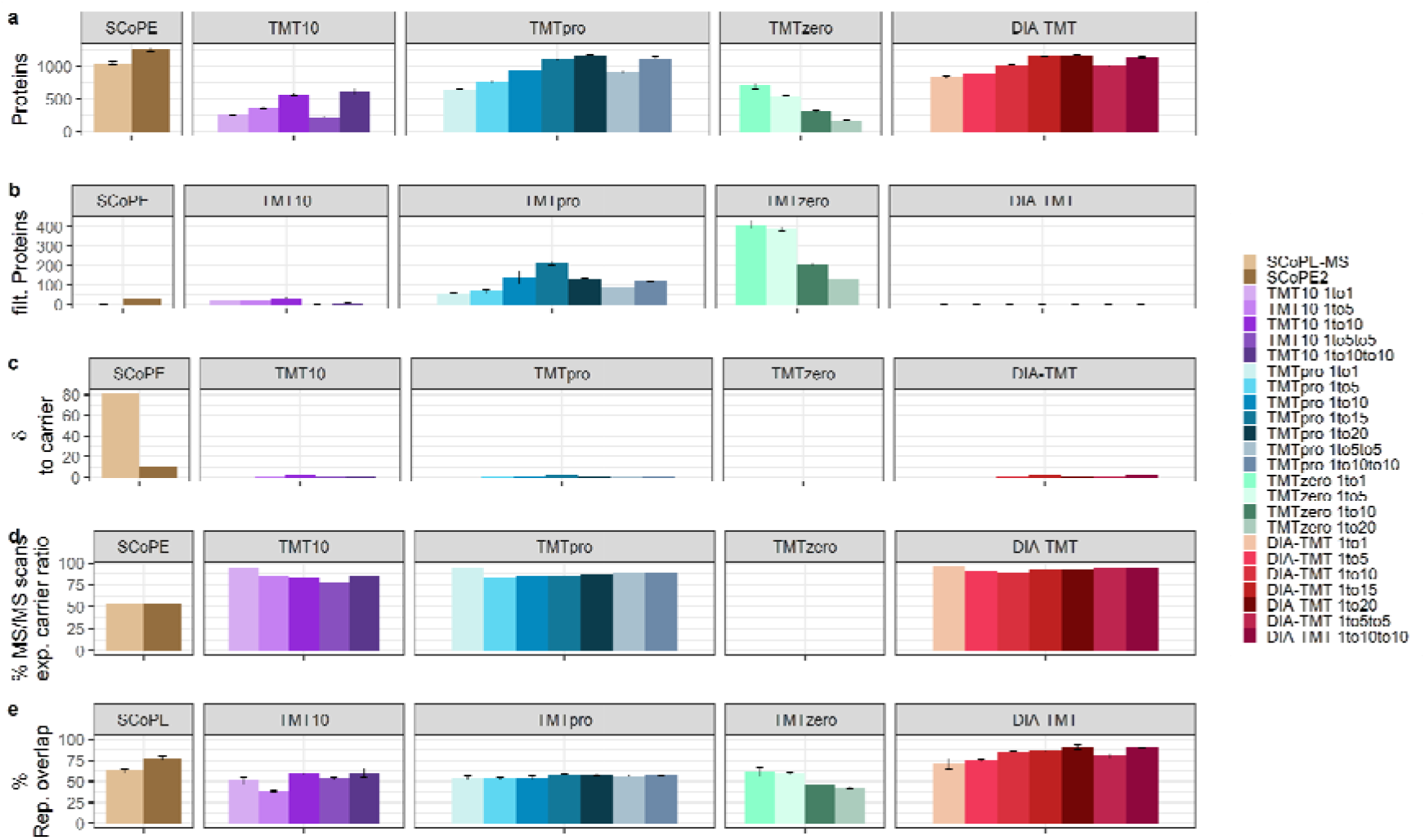
Cumulative comparison of measurement accuracy, variance, and reproducibility across all experimental setups. **(a)** Identified proteins groups, **(b)** S/N filtered protein groups (of note: DIA-TMTpro panel is not included as single MS/MS scans do not give rise to protein identifications), **(c)** delta carrier to ‘single cell’ intensities (optimal: 0), **(d)** percent of MS/MS scans with ‘single cell’ RI within +/− 50% of expected carrier ratio (of note: TMT_zero_ is not included for c and d as the ‘single cell’ carrier ratio cannot be determined within one MS/MS scan) and **(e)** percent replicate overlap across triplicates based on unique peptides for SCoPE (brown), TMT10 (purple), TMTpro (blue), TMT_zero_ (turquoise) and DIA-TMT (red). Bar graphs display median and error bars indicate mad.

Moreover, the no-carrier samples demonstrated outstanding measurement accuracy and reduced variability, especially in TMTpro samples (Fig. 2c, e; Fig. 5c-d; Supplemental Fig. 3c, e). Despite comparable peptide input and total ion current of 10× TMT10 or 5× TMTpro samples, the latter yielded 25% more MS/MS scans and protein identifications. Due to similar ‘single cell’ CVs with both isobaric reagents, we speculate that the global identification-increase with TMTpro results from different fragmentation patterns (Fig. 1b). This further provided a major advantage to overcome detrimental precursor stochasticity and improve reproducibility with our multiplexed DIA strategy. Consequentially, the DIA-TMT acquisition of TMTpro samples provided comparable protein identifications, superior data completeness resulting in close to 100% replicate overlap at reduced measurement variance (Fig. 4b-h; Fig. 5a-e).

The alternative triggering via TMT_zero_ confirmed that abundant carrier spikes dominate low abundant ‘single cell’ MS/MS spectra, even if segregated from the RI cluster (Fig. 3c-d). Importantly, such co-isolated MS/MS spectra may comprise mainly of carrier b- and y-ions while the presence of any RI signal is used for quantification of ‘single cells’.^13,14^ Even though high TMT_zero_ RI intensities indicate that the acquisition strategy might overestimate the prevalence of mixed spectra, already at ≥5x carrier spikes most MS/MS scans either comprise only carrier or co-isolated precursors (Fig. 3d; Supplemental Fig. 5a-c). While background and possible contaminations are reduced to a minimum in our bulk dilutions, this might affect the biological interpretation of real single-cell samples. Contrarily, the intentional co-isolation in a prescheduled acquisition scheme using DIA-TMT successfully defines cell-types, underrepresented single protein knockouts, and presents less quantitative variance at theoretically no missing data.^24^ The need for quantitative data imputation in standard single cell DDA data is highly elevated compared to standard input, while the absence of technical replicates challenges reliability.^12,21,38–40^ Despite the intentional co-isolation in DIA-TMT, eliminating precursor stochasticity drastically improved accuracy and sensitivity (Fig. 4b-f).^24^ Hence we speculate that large proportions computationally generated quantitative data introduced by reduced replicate overlap in combination with precursor co-isolation and extreme carrier ratios are particularly error prone.

We present a comprehensive overview of currently available multiplexed single cell proteomics setups considering protein identifications, measurement variance, quantitative accuracy, and missing data. We find that specific experimental questions require individual prioritization of parameters when designing ultra-low input or single cell studies. Based on these findings, we conclude that limiting carrier spikes (i.e., ≤20x) is pivotal for accurate single-cell proteomics analysis and thus any biological interpretation (Fig. 5a-e). With more sensitive instrumentation and dedicated experimental approaches single-cell proteomics has achieved remarkable proteome depth and throughput. Nevertheless, many parameters such as cell state, sample preparation, chromatography and ultimately the acquisition style impact data quality. We are confident that efficient sample preparation workflows, novel instrumentation and tightly controlled computational approaches will drive biological applications and further demonstrate the impact of hypothesis-free proteome measurements at single-cell resolution.

## Supporting information

Additional experimental details including in-depth ratio suppression and measurement variance for all conditions contained in the manuscript, comparative reanalysis of the published SCoPE datasets, and measurement accuracy of TMT_zero_.

## Acknowledgments

We thank all members of our laboratories for helpful discussions. TMT_zero_ and DIA-TMT were conceptualized together with Johannes Stadlmann, whose support on study design and interpretation was essential. We specifically want to thank Manuel Matzinger and Elisabeth Roitinger for critical comments on the manuscript. This work has been supported by EPIC-XS, project number 823839, funded by the Horizon 2020 program of the European Union and the Austrian Science Fund by ERA-CAPS I 3686-B25-MEIOREC international project.

## Data Availability

All mass spectrometry-based proteomics data have been deposited to the ProteomeXchange Consortium via the PRIDE partner repository with the dataset identifier [PXD027912]

Reviewer account details: Username, Password: G1jNd1XX

## References

(1) Zhu, Y.; Scheibinger, M.; Ellwanger, D. C.; Krey, J. F.; Choi, D.; Kelly, R. T.; Heller, S.; Barr-Gillespie, P. G. Single-Cell Proteomics Reveals Changes in Expression during Hair-Cell Development. eLife 2019, 8, e50777. https://doi.org/10.7554/eLife.50777.

(2) Izar, B.; Tirosh, I.; Stover, E. H.; Wakiro, I.; Cuoco, M. S.; Alter, I.; Rodman, C.; Leeson, R.; Su, M.-J.; Shah, P.; Iwanicki, M.; Walker, S. R.; Kanodia, A.; Melms, J. C.; Mei, S.; Lin, J.-R.; Porter, C. B. M.; Slyper, M.; Waldman, J.; Jerby-Arnon, L.; Ashenberg, O.; Brinker, T. J.; Mills, C.; Rogava, M.; Vigneau, S.; Sorger, P. K.; Garraway, L. A.; Konstantinopoulos, P. A.; Liu, J. F.; Matulonis, U.; Johnson, B. E.; Rozenblatt-Rosen, O.; Rotem, A.; Regev, A. A Single-Cell Landscape of High-Grade Serous Ovarian Cancer. Nat. Med. 2020, 26 (8), 1271–1279. https://doi.org/10.1038/s41591-020-0926-0.

(3) Lombard‐Banek, C.; Moody, S. A.; Nemes, P. Single-Cell Mass Spectrometry for Discovery Proteomics: Quantifying Translational Cell Heterogeneity in the 16-Cell Frog (Xenopus) Embryo. Angew. Chem. Int. Ed. 2016, 55 (7), 2454–2458. https://doi.org/10.1002/anie.201510411.

(4) Thul, P. J.; Åkesson, L.; Wiking, M.; Mahdessian, D.; Geladaki, A.; Blal, H. A.; Alm, T.; Asplund, A.; Björk, L.; Breckels, L. M.; Bäckström, A.; Danielsson, F.; Fagerberg, L.; Fall, J.; Gatto, L.; Gnann, C.; Hober, S.; Hjelmare, M.; Johansson, F.; Lee, S.; Lindskog, C.; Mulder, J.; Mulvey, C. M.; Nilsson, P.; Oksvold, P.; Rockberg, J.; Schutten, R.; Schwenk, J. M.; Sivertsson, Å.; Sjöstedt, E.; Skogs, M.; Stadler, C.; Sullivan, D. P.; Tegel, H.; Winsnes, C.; Zhang, C.; Zwahlen, M.; Mardinoglu, A.; Pontén, F.; Feilitzen, K. von; Lilley, K. S.; Uhlén, M.; Lundberg, E. A Subcellular Map of the Human Proteome. Science 2017, 356 (6340). https://doi.org/10.1126/science.aal3321.

(5) Cong, Y.; Motamedchaboki, K.; Misal, S. A.; Liang, Y.; Guise, A. J.; Truong, T.; Huguet, R.; Plowey, E. D.; Zhu, Y.; Lopez-Ferrer, D.; Kelly, R. T. Ultrasensitive Single-Cell Proteomics Workflow Identifies >1000 Protein Groups per Mammalian Cell. Chem. Sci. 2021. https://doi.org/10.1039/D0SC03636F.

(6) Brunner, A.-D.; Thielert, M.; Vasilopoulou, C.; Ammar, C.; Coscia, F.; Mund, A.; Horning, O. B.; Bache, N.; Apalategui, A.; Lubeck, M.; Raether, O.; Park, M. A.; Richter, S.; Fischer, D. S.; Theis, F. J.; Meier, F.; Mann, M. Ultra-High Sensitivity Mass Spectrometry Quantifies Single-Cell Proteome Changes upon Perturbation. bioRxiv 2020, 2020.12.22.423933. https://doi.org/10.1101/2020.12.22.423933.

(7) Specht, H.; Emmott, E.; Petelski, A. A.; Huffman, R. G.; Perlman, D. H.; Serra, M.; Kharchenko, P.; Koller, A.; Slavov, N. Single-Cell Proteomic and Transcriptomic Analysis of Macrophage Heterogeneity Using SCoPE2. Genome Biol. 2021, 22 (1), 50. https://doi.org/10.1186/s13059-021-02267-5.

(8) Budnik, B.; Levy, E.; Harmange, G.; Slavov, N. SCoPE-MS: Mass Spectrometry of Single Mammalian Cells Quantifies Proteome Heterogeneity during Cell Differentiation. Genome Biol. 2018, 19 (1). https://doi.org/10.1186/s13059-018-1547-5.

(9) Dou, M.; Clair, G.; Tsai, C.-F.; Xu, K.; Chrisler, W. B.; Sontag, R. L.; Zhao, R.; Moore, R. J.; Liu, T.; Pasa-Tolic, L.; Smith, R. D.; Shi, T.; Adkins, J. N.; Qian, W.-J.; Kelly, R. T.; Ansong, C.; Zhu, Y. High-Throughput Single Cell Proteomics Enabled by Multiplex Isobaric Labeling in a Nanodroplet Sample Preparation Platform. Anal. Chem. 2019, 91 (20), 13119–13127. https://doi.org/10.1021/acs.analchem.9b03349.

(10) Schoof, E. M.; Furtwängler, B.; Üresin, N.; Rapin, N.; Savickas, S.; Gentil, C.; Lechman, E.; Keller, U. auf dem; Dick, J. E.; Porse, B. T. Quantitative Single-Cell Proteomics as a Tool to Characterize Cellular Hierarchies. Nat. Commun. 2021, 12 (1), 3341. https://doi.org/10.1038/s41467-021-23667-y.

(11) Schoof, E. M.; Rapin, N.; Savickas, S.; Gentil, C.; Lechman, E.; Haile, J. S.; Keller, U. auf dem; Dick, J. E.; Porse, B. T. A Quantitative Single-Cell Proteomics Approach to Characterize an Acute Myeloid Leukemia Hierarchy. bioRxiv 2019, 745679. https://doi.org/10.1101/745679.

(12) Specht, H.; Slavov, N. Optimizing Accuracy and Depth of Protein Quantification in Experiments Using Isobaric Carriers. J. Proteome Res. 2020. https://doi.org/10.1021/acs.jproteome.0c00675.

(13) Cheung, T. K.; Lee, C.-Y.; Bayer, F. P.; McCoy, A.; Kuster, B.; Rose, C. M. Defining the Carrier Proteome Limit for Single-Cell Proteomics. Nat. Methods 2020, 1–8. https://doi.org/10.1038/s41592-020-01002-5.

(14) Stopfer, L. E.; Conage-Pough, J. E.; White, F. M. Quantitative Consequences of Protein Carriers in Immunopeptidomics and Tyrosine Phosphorylation MS2 Analyses. Mol. Cell. Proteomics 2021, 20. https://doi.org/10.1016/j.mcpro.2021.100104.

(15) O’Brien, J. J.; O’Connell, J. D.; Paulo, J. A.; Thakurta, S.; Rose, C. M.; Weekes, M. P.; Huttlin, E. L.; Gygi, S. P. Compositional Proteomics: Effects of Spatial Constraints on Protein Quantification Utilizing Isobaric Tags. J. Proteome Res. 2018, 17 (1), 590–599. https://doi.org/10.1021/acs.jproteome.7b00699.

(16) Kelstrup, C. D.; Aizikov, K.; Batth, T. S.; Kreutzman, A.; Grinfeld, D.; Lange, O.; Mourad, D.; Makarov, A. A.; Olsen, J. V. Limits for Resolving Isobaric Tandem Mass Tag Reporter Ions Using Phase-Constrained Spectrum Deconvolution. J. Proteome Res. 2018. https://doi.org/10.1021/acs.jproteome.8b00381.

(17) Savitski, M. M.; Mathieson, T.; Zinn, N.; Sweetman, G.; Doce, C.; Becher, I.; Pachl, F.; Kuster, B.; Bantscheff, M. Measuring and Managing Ratio Compression for Accurate ITRAQ/TMT Quantification. J. Proteome Res. 2013, 12 (8), 3586–3598. https://doi.org/10.1021/pr400098r.

(18) Tsai, C.-F.; Zhao, R.; Williams, S. M.; Moore, R. J.; Schultz, K.; Chrisler, W. B.; Pasa-Tolic, L.; Rodland, K. D.; Smith, R. D.; Shi, T.; Zhu, Y.; Liu, T. An Improved Boosting to Amplify Signal with Isobaric Labeling (IBASIL) Strategy for Precise Quantitative Single-Cell Proteomics. Mol. Cell. Proteomics 2020, 19 (5), 828–838. https://doi.org/10.1074/mcp.RA119.001857.

(19) Furtwängler, B.; Üresin, N.; Motamedchaboki, K.; Huguet, R.; Lopez-Ferrer, D.; Zabrouskov, V.; Porse, B. T.; Schoof, E. M. Real-Time Search Assisted Acquisition on a Tribrid Mass Spectrometer Improves Coverage in Multiplexed Single-Cell Proteomics; 2021; p 2021.08.16.456445. https://doi.org/10.1101/2021.08.16.456445.

(20) Peshkin, L.; Gupta, M.; Ryazanova, L.; Wühr, M. Bayesian Confidence Intervals for Multiplexed Proteomics Integrate Ion-Statistics with Peptide Quantification Concordance*, [S]. Mol. Cell. Proteomics 2019, 18 (10), 2108–2120. https://doi.org/10.1074/mcp.TIR119.001317.

(21) Brenes, A.; Hukelmann, J.; Bensaddek, D.; Lamond, A. I. Multibatch TMT Reveals False Positives, Batch Effects and Missing Values. Mol. Cell. Proteomics 2019, 18 (10), 1967–1980. https://doi.org/10.1074/mcp.RA119.001472.

(22) Kovalchik, K. A.; Colborne, S.; Spencer, S. E.; Sorensen, P. H.; Chen, D. D. Y.; Morin, G. B.; Hughes, C. S. RawTools: Rapid and Dynamic Interrogation of Orbitrap Data Files for Mass Spectrometer System Management. J. Proteome Res. 2019, 18 (2), 700–708. https://doi.org/10.1021/acs.jproteome.8b00721.

(23) Hulsen, T.; de Vlieg, J.; Alkema, W. BioVenn – a Web Application for the Comparison and Visualization of Biological Lists Using Area-Proportional Venn Diagrams. BMC Genomics 2008, 9 (1), 488. https://doi.org/10.1186/1471-2164-9-488.

(24) Ctortecka, C.; Krššáková, G.; Stejskal, K.; Penninger, J. M.; Mendjan, S.; Mechtler, K.; Stadlmann, J. Comparative Proteome Signatures of Trace Samples by Multiplexed Data-Independent Acquisition. Mol. Cell. Proteomics 2021, 0 (0). https://doi.org/10.1016/j.mcpro.2021.100177.

(25) Ting, L.; Rad, R.; Gygi, S. P.; Haas, W. MS3 Eliminates Ratio Distortion in Isobaric Multiplexed Quantitative Proteomics. Nat. Methods 2011, 8 (11), 937–940. https://doi.org/10.1038/nmeth.1714.

(26) Savitski, M. M.; Sweetman, G.; Askenazi, M.; Marto, J. A.; Lang, M.; Zinn, N.; Bantscheff, M. Delayed Fragmentation and Optimized Isolation Width Settings for Improvement of Protein Identification and Accuracy of Isobaric Mass Tag Quantification on Orbitrap-Type Mass Spectrometers. Anal. Chem. 2011, 83 (23), 8959–8967. https://doi.org/10.1021/ac201760x.

(27) Paulo, J. A.; O’Connell, J. D.; Gygi, S. P. A Triple Knockout (TKO) Proteomics Standard for Diagnosing Ion Interference in Isobaric Labeling Experiments. J. Am. Soc. Mass Spectrom. 2016, 27 (10), 1620–1625. https://doi.org/10.1007/s13361-016-1434-9.

(28) Paulo, J. A.; Navarrete-Perea, J.; Guha Thakurta, S.; Gygi, S. P. TKO6: A Peptide Standard to Assess Interference for Unit-Resolved Isobaric Labeling Platforms. J. Proteome Res. 2018. https://doi.org/10.1021/acs.jproteome.8b00902.

(29) Navarrete-Perea, J.; Gygi, S. P.; Paulo, J. A. HYpro16: A Two-Proteome Mixture to Assess Interference in Isobaric Tag-Based Sample Multiplexing Experiments. J. Am. Soc. Mass Spectrom. 2020. https://doi.org/10.1021/jasms.0c00299.

(30) Virreira Winter, S.; Meier, F.; Wichmann, C.; Cox, J.; Mann, M.; Meissner, F. EASI-Tag Enables Accurate Multiplexed and Interference-Free MS2-Based Proteome Quantification. Nat. Methods 2018, 15 (7), 527–530. https://doi.org/10.1038/s41592-018-0037-8.

(31) Karp, N. A.; Huber, W.; Sadowski, P. G.; Charles, P. D.; Hester, S. V.; Lilley, K. S. Addressing Accuracy and Precision Issues in ITRAQ Quantitation. Mol. Cell. Proteomics MCP 2010, 9 (9), 1885–1897. https://doi.org/10.1074/mcp.M900628-MCP200.

(32) Lazar, C.; Gatto, L.; Ferro, M.; Bruley, C.; Burger, T. Accounting for the Multiple Natures of Missing Values in Label-Free Quantitative Proteomics Data Sets to Compare Imputation Strategies. J. Proteome Res. 2016, 15 (4), 1116–1125. https://doi.org/10.1021/acs.jproteome.5b00981.

(33) O’Brien, J. J.; Gunawardena, H. P.; Paulo, J. A.; Chen, X.; Ibrahim, J. G.; Gygi, S. P.; Qaqish, B. F. The Effects of Nonignorable Missing Data on Label-Free Mass Spectrometry Proteomics Experiments. Ann. Appl. Stat. 2018, 12 (4), 2075–2095. https://doi.org/10.1214/18-AOAS1144.

(34) Karpievitch, Y. V.; Dabney, A. R.; Smith, R. D. Normalization and Missing Value Imputation for Label-Free LC-MS Analysis. BMC Bioinformatics 2012, 13 (16), S5. https://doi.org/10.1186/1471-2105-13-S16-S5.

(35) Hartlmayr, D.; Ctortecka, C.; Seth, A.; Mendjan, S.; Tourniaire, G.; Mechtler, K. An Automated Workflow for Label-Free and Multiplexed Single Cell Proteomics Sample Preparation at Unprecedented Sensitivity. bioRxiv 2021, 2021.04.14.439828. https://doi.org/10.1101/2021.04.14.439828.

(36) Erickson, B. K.; Rose, C. M.; Braun, C. R.; Erickson, A. R.; Knott, J.; McAlister, G. C.; Wühr, M.; Paulo, J. A.; Everley, R. A.; Gygi, S. P. A Strategy to Combine Sample Multiplexing with Targeted Proteomics Assays for High-Throughput Protein Signature Characterization. Mol. Cell 2017, 65 (2), 361–370. https://doi.org/10.1016/j.molcel.2016.12.005.

(37) Schweppe, D. K.; Eng, J. K.; Yu, Q.; Bailey, D.; Rad, R.; Navarrete-Perea, J.; Huttlin, E. L.; Erickson, B. K.; Paulo, J. A.; Gygi, S. P. Full-Featured, Real-Time Database Searching Platform Enables Fast and Accurate Multiplexed Quantitative Proteomics. J. Proteome Res. 2020, 19 (5), 2026–2034. https://doi.org/10.1021/acs.jproteome.9b00860.

(38) Vanderaa, C.; Gatto, L. Utilizing Scp for the Analysis and Replication of Single-Cell Proteomics Data. bioRxiv 2021, 2021.04.12.439408. https://doi.org/10.1101/2021.04.12.439408.

(39) Lim, M. Y.; Paulo, J. A.; Gygi, S. P. Evaluating False Transfer Rates from the Match-Between-Runs Algorithm with a Two-Proteome Model. J. Proteome Res. 2019. https://doi.org/10.1021/acs.jproteome.9b00492.

(40) Yu, F.; Haynes, S. E.; Nesvizhskii, A. I. IonQuant Enables Accurate and Sensitive Label-Free Quantification With FDR-Controlled Match-Between-Runs. Mol. Cell. Proteomics 2021, 20, 100077. https://doi.org/10.1016/j.mcpro.2021.100077.

