## Supplemental Figures for "Quantitative accuracy and precision in multiplexed single-cell proteomics"

### Affiliations:

### Table of Content:

Supplemental Figure 1: *Re-analysis of public SCoPE data.*

Supplemental Figure 2: *Ratio compression for SCoPE, TMT10-plex and TMTpro samples at various carrier spikes.*

Supplemental Figure 3: *Measurement stability, RI variability and quantification accuracy at various carrier ratios.*

Supplemental Figure 4: *Measurement stability and variance of public SCoPE data.*

Supplemental Figure 5: *Measurement stability and accuracy of TMT_zero_ experiments.*

Supplemental Figure 6: *Ratio compression in DIA-TMT experiments.*

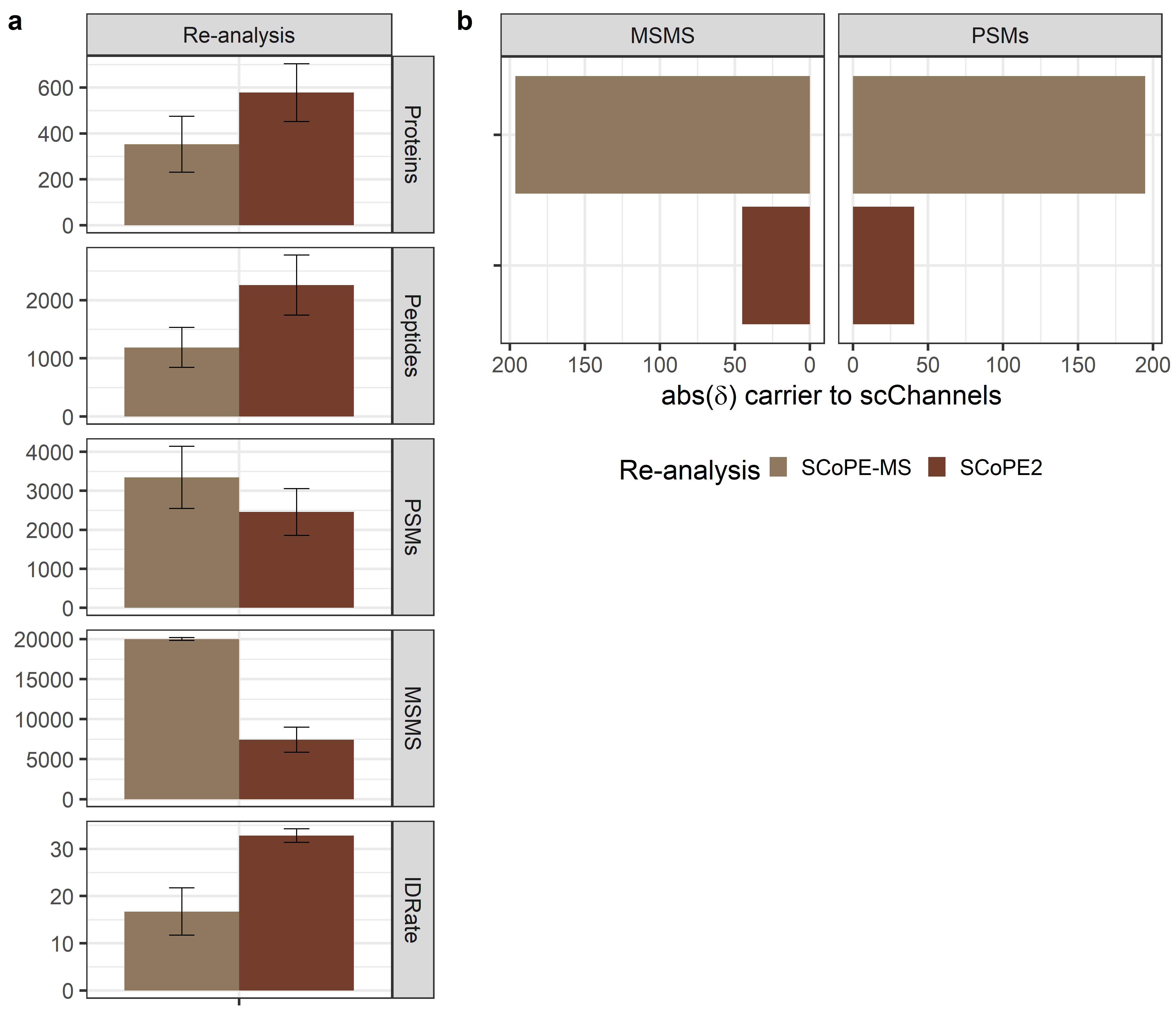
*Supplemental Figure 1: Re-analysis of public single cell data,* **(a)** Identified proteins, peptide groups, PSMs, number of MS/MS scans, ID-rates and delta between expected and acquired carrier to single cell ratio across all MS/MS scans or PSMs of published SCoPE-MS (light brown) and SCoPE2 (dark brown). Median and median absolute deviation (mad) is shown.

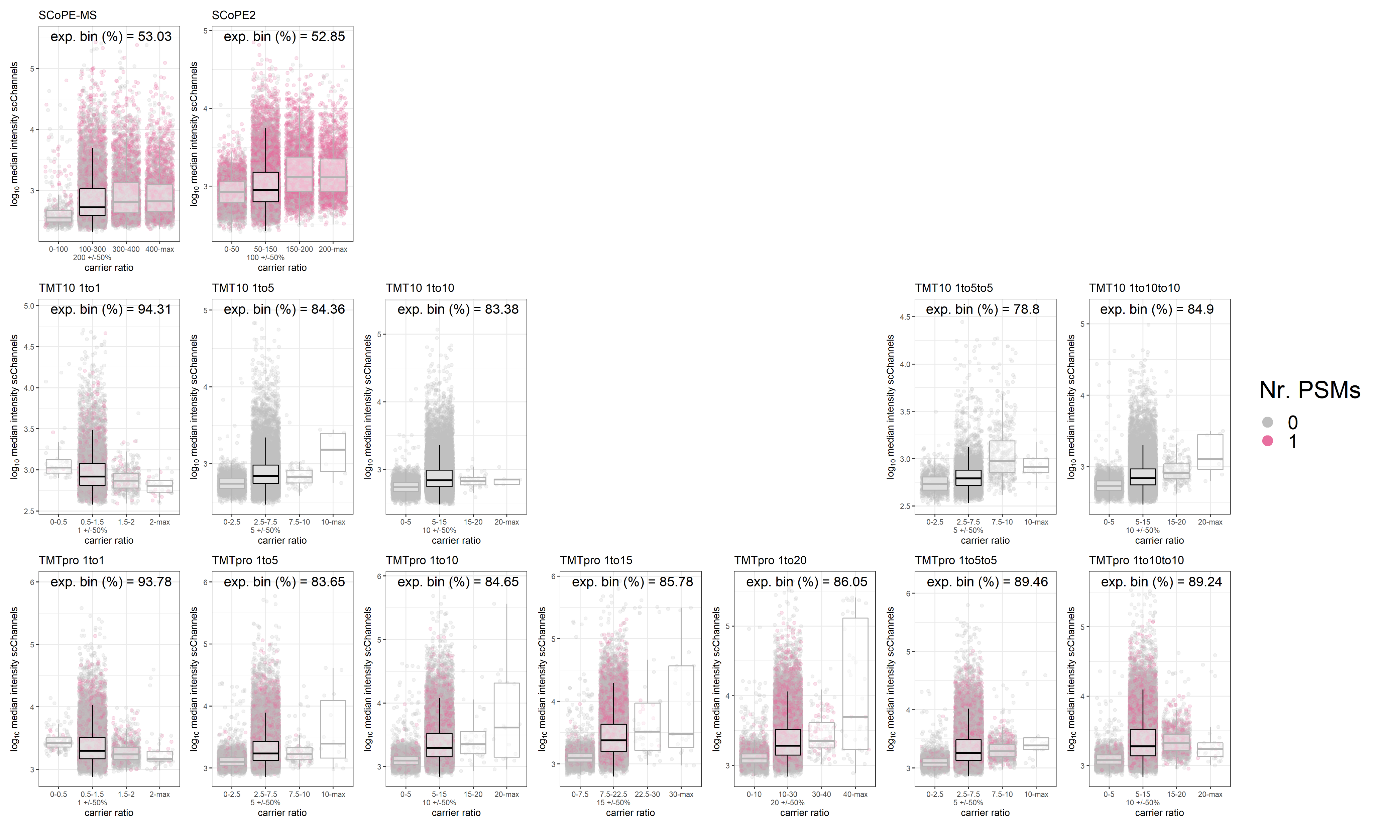

*Supplemental Figure 2: Ratio compression for SCoPE, TMT10-plex and TMTpro samples at various carrier spikes.* Log_10_ median RI intensity of all MS/MS scans and binned ratios between ‘single cells’ and the carrier is displayed. Identified and unidentified MS/MS scans are indicated in grey or pink, respectively. The expected bin (carrier +/- 50 %) is highlighted in black, and the percent of MS/MS scans within those is indicated.

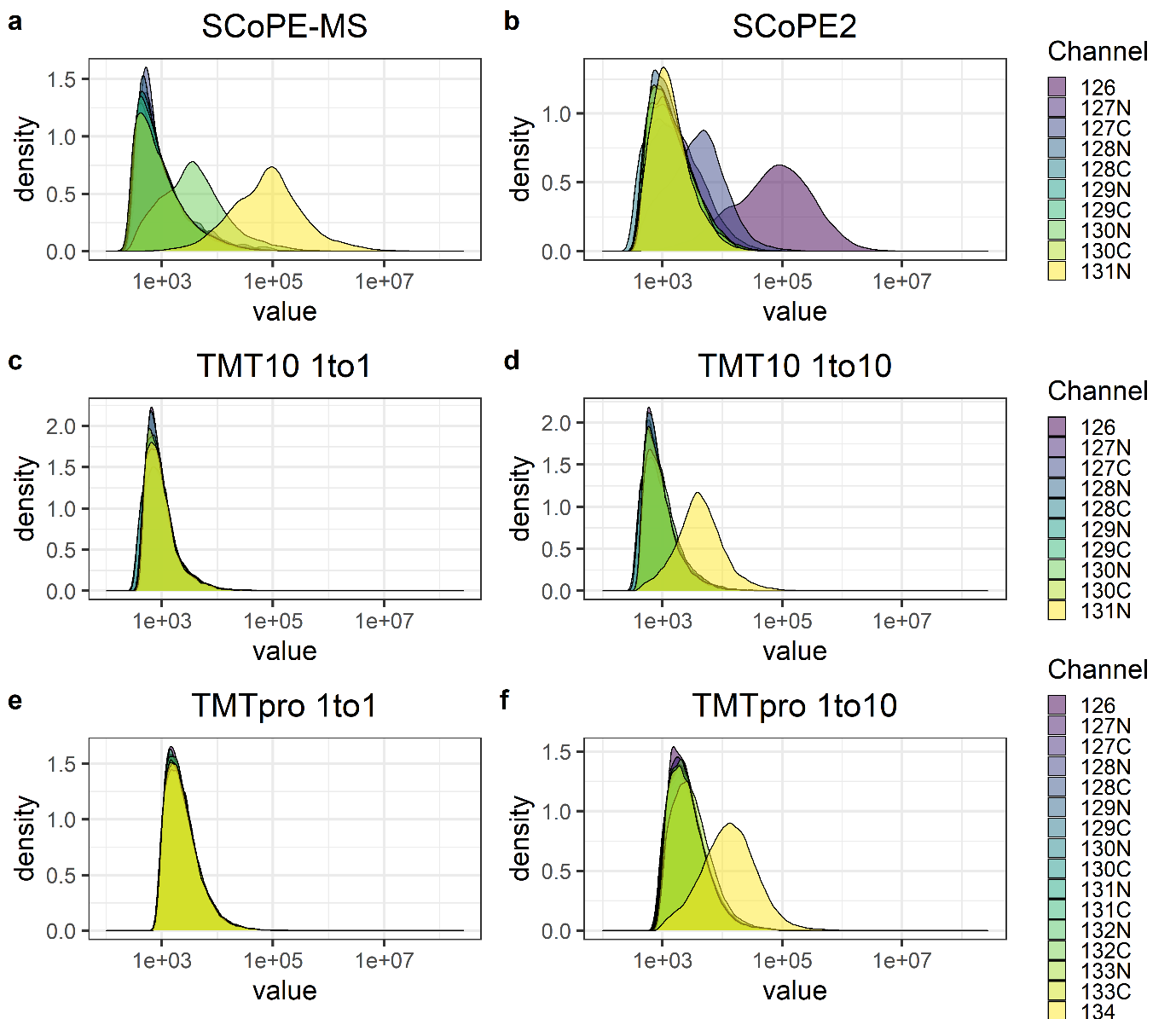

*Supplemental Figure 3: Measurement stability and RI variability at various carrier ratios.* RI intensity distributions based on all MS/MS scans for **(a-b)** SCoPE, **(c-d)** TMT10-plex and **(e-f)** TMTpro experiments at indicated carrier spikes.

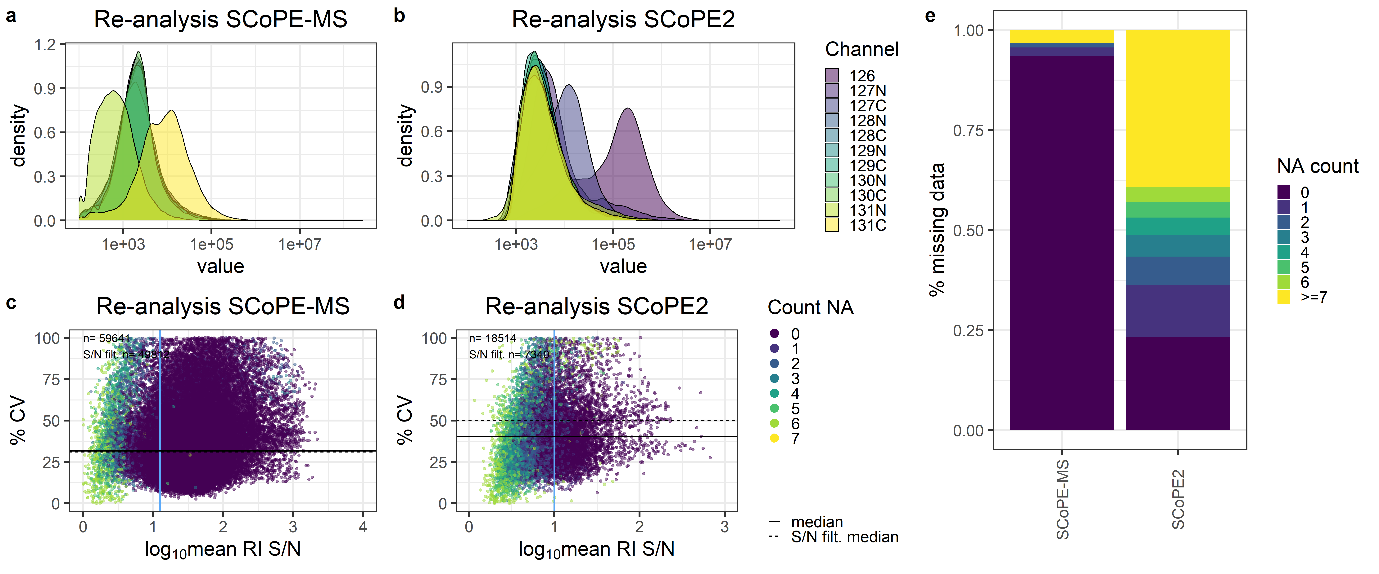
*Supplemental Figure 4: Measurement stability and variance of public SCoPE-MS and SCoPE2 data.* **(a-b)** RI intensity distributions based on all MS/MS scans and **(c-d)** percent CV across ‘single cell’ channels and log_10_ mean RI S/N for publicly available SCoPE data. The horizontal solid line and dashed lines indicate median S/N across all MS/MS scans or post-S/N filtering, respectively. The vertical blue line specifies the S/N filter cut-off. Colors reflect the number of missing ‘single cell’ RIs per MS/MS scan. **(e)** Percent missing quantitative data in publicly available SCoPE data per PSM.

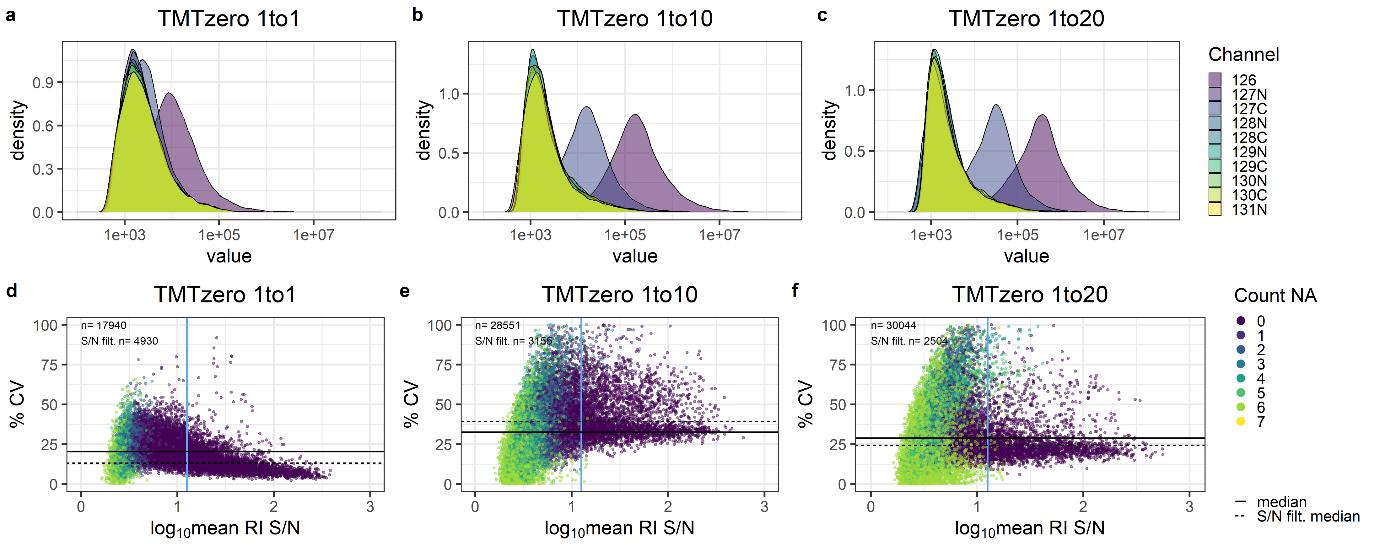

*Supplemental Figure 5: Measurement accuracy of TMT_zero_ experiments*. **(a-c)** RI intensity distribution across all MS/MS scans in TMT_zero_ experiments. Colors indicate different TMT channels. **(d-f)** Median percent CV and log_10_ mean RI S/N at indicated TMT_zero_ carrier spikes. Horizontal solid and dashed lines indicate median S/N across all MS/MS scans or post-S/N filtering, respectively. The vertical blue line indicates the S/N filter cut-off. Colors indicate the number of missing ‘single cell’ RIs per MS/MS scan.

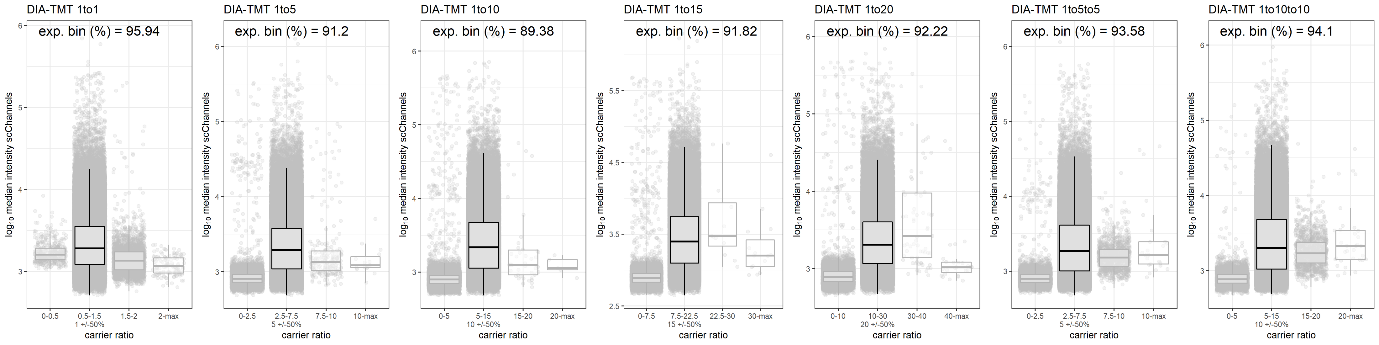

*Supplemental Figure 6: Ratio compression in DIA-TMT experiments.* Log_10_ median RI intensity across all MS/MS scans is displayed with binned ‘single cell’ to carrier channels ratios. The expected ratio (carrier +/- 50%) is highlighted in black, and percent of MS/MS scans within this bin is indicated.
